# PandoraRL: DQN and Graph Convolution based ligand pose learning for SARS-COV1 Mprotease

**DOI:** 10.1101/2022.06.09.495578

**Authors:** Justin Jose, Ujjaini Alam, Pooja Arora, Divye Singh, Nidhi Jatana

## Abstract

The ability to predict the correct ligand binding pose for proteinligand complex is vital for drug design. Recently several machine learning methods have suggested knowledge based scoring functions for binding energy prediction. In this study, we propose a reinforcement learning (RL) based model, PandoraRL, where the RL agent helps the ligand traverse to the optimal binding pose. The underlying representation of molecules utilizes generalized graph convolution to represent the protein ligand complex with various atomic and spatial features. The representation consists of edges formed on the basis of inter molecular interactions such as hydrogen bonds, hydrophobic interactions, etc, and nodes representing atomic features. This study presents our initial model which can train on a protein-ligand pair and predict optimal binding pose for a different ligand with the same protein. To the best of our knowledge, this is the first time an RL based approach has been put forward for predicting optimized ligand pose.

**CCS CONCEPTS:**

- **Computing methodologies** → **Reinforcement learning**.

## 1 INTRODUCTION

Molecular docking is one of the most popular and successful strategies to identify the binding ligand pose, orientation and conformation [19]. The basic theories of docking includes sampling algorithms and scoring functions. Various docking method are reviewed in [15], highlighting their success in ligand pose identification. In spite of their invaluable contributions, a common challenge for these methods is that often the minimal binding energy ligand pose provided by these methods may not be the biologically relevant pose [4]. This has led to the use of machine learning and AI driven methods for predicting binding affinity and ligand conformations, and effective representation of the molecules becomes crucial. Graph based methods have gained popularity due to their ability to provide learnable feature vectors (fingerprints) of defined length even for molecules of varying size [1, 2, 6, 7, 25]. These methods represent molecules as 2D graphs. Many methods have been introduced which utilize graphical representation. Shen et. al. proposed a cascade convolution calculation method for graph embedded representation of protein-ligand pair [23]. In [14], graph and mesh representation was used for the molecules to predict the binding conformation. Recently, [9] and [24] used k-NN graph representation with SE(3)-equivariant geometric deep learning model to tackle the protein-ligand binding problem.

Reinforcement learning (RL), a known technique in the field of optimization, has also made some recent progress in the protein structure world. It is predominantly used as a supplementary aid in generative drug design [28]. In [21] the authors explored protein-ligand docking problem as a one step optimization problem with the molecule moving in one dimension. Jose et. al. proposed a method to represent protein and ligand as a graph and use 3D CNN-based RL algorithm for the rigid protein-ligand docking problem [11]. Recently, [27] used voxel-based representation, with a single atom ligand (copper ion) considered for the simulation process. The protein-ligand docking problem is different from any other RL problem where either the path or the final destination is known. Here each of the protein-ligand interactions differs in– 1. The position of catalytic diad/triad; 2. The type of interactions they form; 3. Shape and nature of the interacting ligand. This makes it complex for the RL agent to learn the diversity each protein ligand pair brings in and to create a model for unseen protein-ligand complexes.

In this study, we put forward a method that employs RL for the identification of optimized ligand pose. The process starts with graphical representation of the protein-ligand complex, which aids the RL algorithm in understanding interactions. This protein-ligand complex graph is then passed on to the graph-based RL algorithm which approximates the underlying molecular interactions using input atomic and spatial features. In our formulation, the RL agent is responsible for identifying the optimized ligand pose.

## 2 MATERIALS AND METHODS

In this section, we elaborate on the dataset used, the data representation, and the network design, which includes the RL process. The overall workflow is shown in Figure 1.

**Figure 1:**
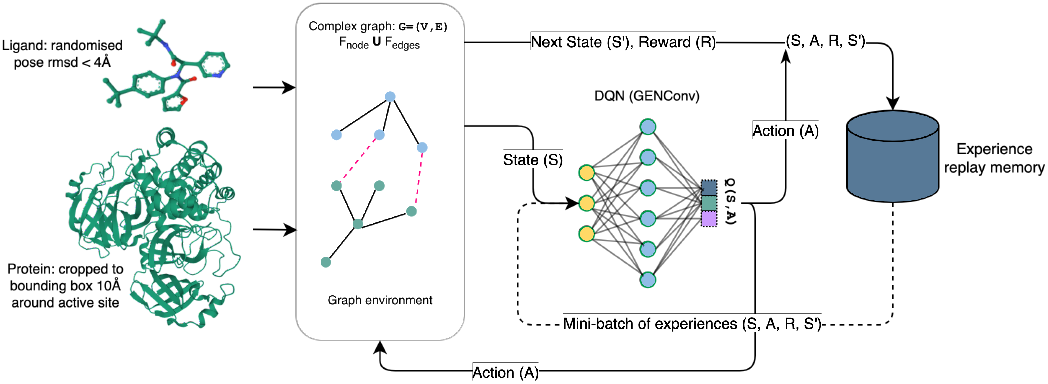
Overall training workflow including protein ligand preparation, graph representation and DQN training.

### 2.1 Dataset

The dataset consists of 11 SARS-Cov1 3C-like main protease (3CL-Pro) protein ligand complexes curated from PDB^1^ and PDBBind^2^, which is randomly split into a group of 8 complexes for training and 3 complexes for testing. To validate the retrain-ability of the RL algorithm, a second set of 7 SARS-COV papain-like protease (PLpro) protein complexes is also utilized.

The complex was split into protein and ligand. The proteins in the complexes are prepared using dock prep [22] tool from Chimera[20], ligands are prepared by adding hydrogen and partial charge and removing water molecules using open babel [18]. The ligand molecule and the receptor molecule are then split into separate files, and the receptor is cropped into a 10Å × 10Å × 10Å cube around the bound ligand position.

### 2.2 Data representation

We represent the protein-ligand complex using a single graph. The protein-ligand graph 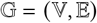 uses atoms as nodes and interactions as the edges. We use open babel [18] to identify intramolecular interactions. We also include the inter-molecular interactions, identified using the thresholds mentioned in [3], [5]. These inter-molecular edges represent the possibility of an interaction, and get updated with every ligand pose change. This enables the reinforcement model to identify the edges that contribute to the optimal pose.

The node feature-set consists of atomic and substructure features. The atomic features include atom type encoding, spatial coordinates, Van-der Waal radius, hybridization, etc. referred to as SMARTS and the residue level features include the residue labels, conformation similarities and amino acid charge for the residues. We replicate these features across all member atoms.

The edge features consist of the one hot encoded bond type (representing covalent, strong and weak hydrogen bond, hydrophobic bond, pi stacking, salt bridge, amide stacking, halogen bond, multipolar halogen bond, cation pi and repulsive bond), the bond energy, and the bond distance.

### 2.3 Network Design

The network design for RL is based on the GENeralized Graph Convolution (GENConv) [12] network. This network design allows edge features to be included along with node features as part of the input data. The network generates an intermediate node level representation through message passing and captures the edge information by summing the edge features with the node features of member nodes of that edge. To obtain an aggregated graph representation, we use average and max pooling layers on the intermediate node representation. The aggregated intermediate graph representation is used by a policy layer, which then updates the pose based on the predicted policy.

One of the major challenges in modelling the ligand pose prediction is that the actions taken to move between different ligand poses are continuous in nature. RL algorithms like Deep Deterministic Policy Gradient (DDPG) [13] work on continuous action space, but suffer from convergence issues [8]. We therefore chose to work with Deep Q-Network (DQN) [16, 17], an off policy reinforcement learning algorithm that uses neural networks to approximate the state-value function in a *Q*-Learning framework, and demonstrates stable learning. Since the *Q*-values learned through DQN map to a discrete set of policies, the ligand pose update has to be discretized. We chose a step sizes of 0.1Å for translation and 0.5° for rotation to map the pose updates into discrete actions, which are linked to the DQN policies through *Q*-values. Detailed description of how DQN updates the *Q*-values is described in the following section.

#### 2.3.1 Deep Q-Network Optimization

The optimal *Q*-value, and the associated loss are given by:

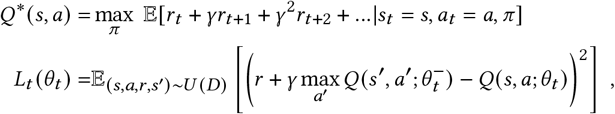

where *Q*\* (*s, a*) is the maximum sum of rewards *r_t_*, discounted by *γ* at each time-step *t*, that can be realized by making an observation s and taking an action *a*, using a behaviour policy *π* = *P*(*α|s*). The *Q*-learning updates are applied on samples (*s, a,r,s*’) ~ *U*(*D*), drawn randomly and uniformly from the stored sample pool. The loss at the *t*-th time-step, *L_t_* (*θ_t_*), is dependent on the parameters of the *Q*-network, *θ_t_*, and the network parameters used to compute the target, 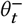, at the *t*-th timestep.

For protein ligand docking, the prediction of an optimal *Q*-value should lead to an optimal approximation of the correct binding pose. In this scenario, the features of the *Q*-network are the node and edge features of the protein-ligand graph described in section 2.2. The optimal *Q*-value depends on the optimization of the reward function, hence the choice of reward function is crucial to the convergence of the algorithm.

#### 2.3.2 Reward function

The reward function has to be chosen such that maximizing it leads the algorithm to the correct pose. It also has to be linked to the features so that the algorithm learns to predict the correct action based on the features at each pose. Minimizing the *rmse* between the ligand pose at any given time and the correct ligand pose naturally trains the algorithm to find the correct pose. A weakness of an *rmse*-based reward function is that it is not explicitly linked to the features of the protein and ligand, but depends only on the distance from the correct pose. However, it lends itself to straightforward optimization, and can therefore be a good starting point. We use a sine hyperbolic reward function of the form ℝ(*x*) = 1/sinh(*rmse^a^*), which decreases monotonically from the value at correct pose, and has no local maxima in the range defined.

#### 2.3.3 Training & Testing

The algorithm is trained iteratively on the SARS-Cov1 training dataset. At each iteration, it picks a random protein-ligand complex, with the ligand at a random pose, and uses translation and rotation actions to move it towards the optimal bound pose, based on the reward function. The ligand is judged sufficiently close to the bound position at an *rmse* = 0.1Å, while an *rmse* = 4Å is too far from the bound pose and is awarded a penalty. This process is repeated for different complexes and starting poses, till the algorithm is able to identify the underlying optimal *Q*-values, and associated actions.

The model thus created is then applied on the test dataset to predict optimal ligand poses. Since *rmse* cannot be used as a stopping condition here, we use Δp, change in pose at each step, to ascertain if a better pose can be generated. If the change in pose is sufficiently small over several steps, and displays oscillatory behaviour, the generated pose is accepted as the optimal ligand pose.

We validate the correctness of the generated optimal ligand-pose by comparing the known bound ligand pose with the optimized ligand pose suggested by the RL model.

## 3 RESULT AND DISCUSSION

In this section we show our results that demonstrate that the RL agent is able to learn protein ligand interactions and to help the ligand traverse towards the favourable pose. To understand the agents learning patterns we have performed experiments with different feature set and reward function combinations as shown in table 1.

**Table 1:** RL training experiments for different feature sets with RMSE as reward function.

| Feature set | outcome |
| --- | --- |
| SMARTS | No learning, ligand oscillates at one place |
| SMARTS + EDGE types | Shows learning in training, starts moving towards protein |
| Coordinates + SMARTS + EDGE types | Shows learning in training, tied to absolute coordinates of each protein ligand complex |
| Distance + SMARTS + EDGE types | Shows learning in training, ties to absolute distances in each protein ligand complex |
| Residue labels + SMARTS + EDGE types | Converging model; moves from RMSE 6Å to 3Å in test and to < 2Å in training |

We infer that the PandoraRL model is able to learn best when supported with features like Residue labels and edges formed of hydrogen bond, hydrophobic interactions, etc. Additionally, the weights of the initial node feature encoding layer identify that atom labels had higher weights and therefore higher impact. Figure 2 shows the model performance for the test data (PDB Id:3v3m). It shows promising results where the ligand placed randomly at a starting position of *rmse* ~ 6Å is able to traverse to *rmse* ~ 3.2Å. We note that the interacting residues for our model as shown in the ligplot^3^ are the same as those for the original bound complex taken from pdbsum^4^. A second complex (PDB id:5C5N) of the SARS-Cov1 test dataset gave similar results, thus showing that RL is able to predict ligand pose to 3Å accuracy for complexes in the with the same protein. We repeated this experiment for the PL-PRO protein-ligand complexes, where the model could minimize *rmse* till ~ 3Å as well. This shows that our framework can be consistently retrained for different protein ligand complex sets with the same protein and different ligands to minimize *rmse* to 3Å accuracy.

**Figure 2:**
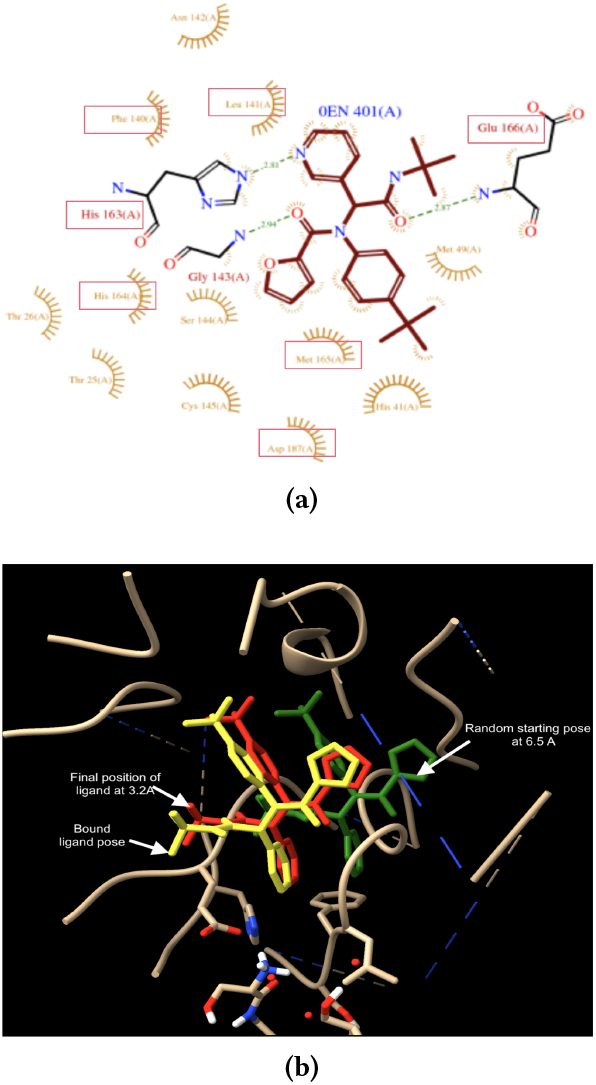
a) Ligplot for 3v3m complex. (b) Comparison of trained ligand pose with the bound ligand. Green represents the starting position at ~ 6Å, red represents the final position at 3.2Å, and yellow represents the original bound ligand.

The current model is able to achieve an RMSE of 3.2Å which can be further optimized to obtain a pose closer to bound ligand. We may improve our current results by extending the current feature set, adding substructure level features such as functional groups for ligands, residue grouping based on charge, hydrophobicity, conformation etc. A comprehensive understanding of the structure and dynamics of the protein-ligand complexes can be obtained by utilizing the NetworkX python package [10] to create the proteinligand graph. We may also experiment with the reward function to link it explicitly to the features. A good candidate for this is Autodock Vina [26], an empirical scoring function that sums up the affinity of protein-ligand binding from different bonds. Another possibility is to calculate the graph similarity between the current and correct poses, which considers the number and types of interactions between protein and ligand atoms. A combination of such feature based reward functions and the *rmse* can provide sufficient information to the algorithm for learning. We expect that these experiments will help the RL algorithm to consistently learn optimized ligand poses, and to predict correct bound poses for a variety of protein ligand complexes to a good degree of accuracy. We plan to generalize our model incrementally by validating it against SARS-Cov2 Mprotease followed by diverse protein ligand complexes.

The above results showcase the potential of RL based methods to learn protein ligand interactions. Given the diversity in protein ligand interactions and their biochemistry, reinforcement learning can make path breaking contributions and tap unknown areas which are less researched due to data scarcity or diversity of data. The result reported above can be regarded as the initial model that validates our hypothesis that RL agent based learning can identify optimized ligand poses. Further refinement is expected to create a more generic algorithm for a variety of data.

## AVAILABILITY

The code for running smaller experiments for reinforcement learning based approach to protein-ligand interaction and the dataset used is available at https://gitlab.com/lifesciences/pandoraRL

## ACKNOWLEDGMENTS

We would like to thank Dr. Manali Joshi, Dr. Jayaraman for interesting discussions and mentoring. We also thank the DDH committee for giving us the opportunity to explore this project.

## Footnotes

1 https://www.rcsb.org/

2 http://www.pdbbind.org.cn/

3 https://www.ebi.ac.uk/thornton-srv/software/LIGPLOT/

4 http://www.ebi.ac.uk/thornton-srv/databases/cgi-bin/pdbsum/GetPage.pl

